## Supplemental Figures and Tables for "*EARLY FLOWERING 3* interactions with *PHYTOCHROME B* and *PHOTOPERIOD1* are critical for the photoperiodic regulation of wheat heading time"

**S1 Fig. Schematic representations of *PHYB*, *ELF3*, and *PPD1* mutants.** Mutations introducing premature stop codons are indicated with red triangles and deletions with dotted red lines. Exons are shown as gray rectangles and introns as black lines. (**A**) The *phyB* line combines a premature stop codon on the A-genome homeolog that truncates the last 641 amino acids of the protein, spanning the entire regulatory module, and a premature stop codon in the B-genome copy that eliminates the distal 140 amino acids, including the histidine kinase domain (Pearce et al., 2016). (**B**) The *elf3* line carries premature stop codons that eliminate the last 241 (A-genome) and 244 (B-genome) amino acids of the C-terminal region, including the third and fourth conserved blocks of the ELF3 protein (Alvarez et al., 2016). (**C**) The *ppd1* line carries a premature stop codon in *ppd-A1* that eliminates 514 amino acids of the PPD-A1 protein, including the highly conserved CCT domain, and *ppd-B1* is a gamma ray-induced deletion the eliminates the complete gene (Pearce et al., 2017; Shaw et al., 2020). (**D**) Natural deletions in the promoter of the *PPD-A1a* (Wilhelm et al., 2009) and *PPD-D1a* (Beales et al., 2006) alleles in tetraploid and hexaploid wheat, respectively.

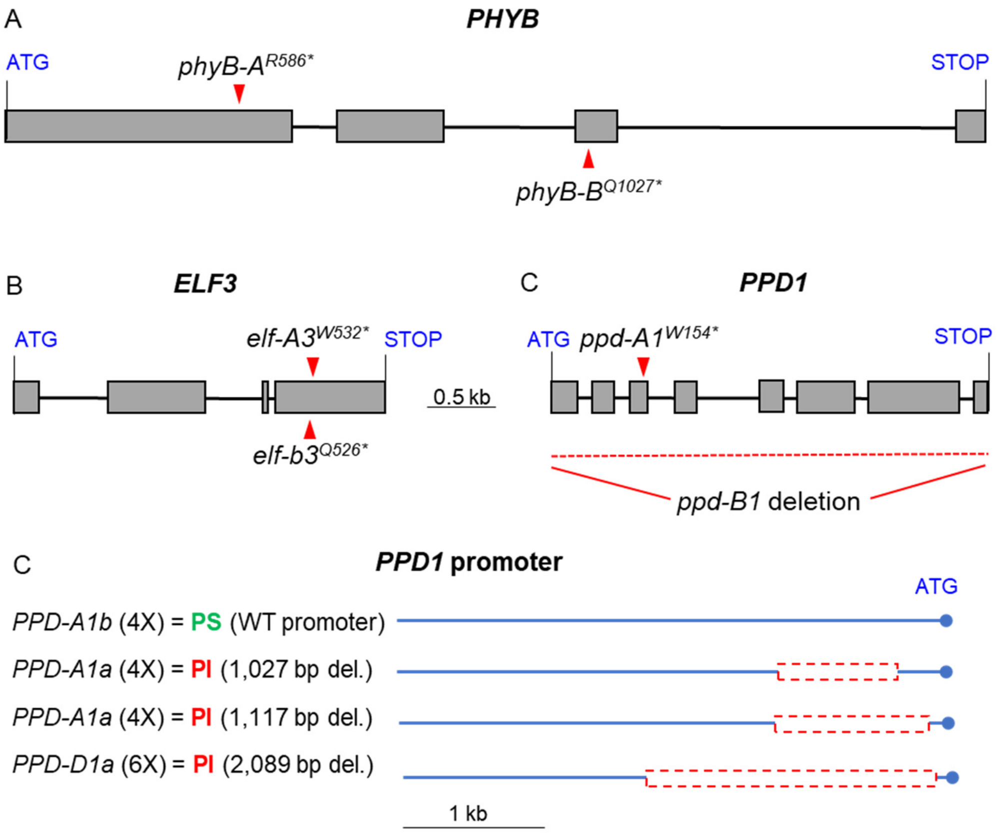

**S2 Fig. Effect of *elf3* and *elf3 phyB* mutations on spikelet number per spike (SNS).** (**A**) Kronos photoperiod insensitive (PI). (**B**) Kronos photoperiod sensitive (PS). In both experiment plants were grown under LD (16h light). Different letters above the bars indicate significant differences in pair-wise non-parametric Kruskal-Wallis tests ($P < 0.05$). The non-parametric test was used because no transformation was able to restore normality of residuals and homogeneity of variances simultaneously. The reduced SNS in the *elf3* mutant was restored in the *phyB elf3* combined mutant. Raw data is available in Data H in S1 Data.

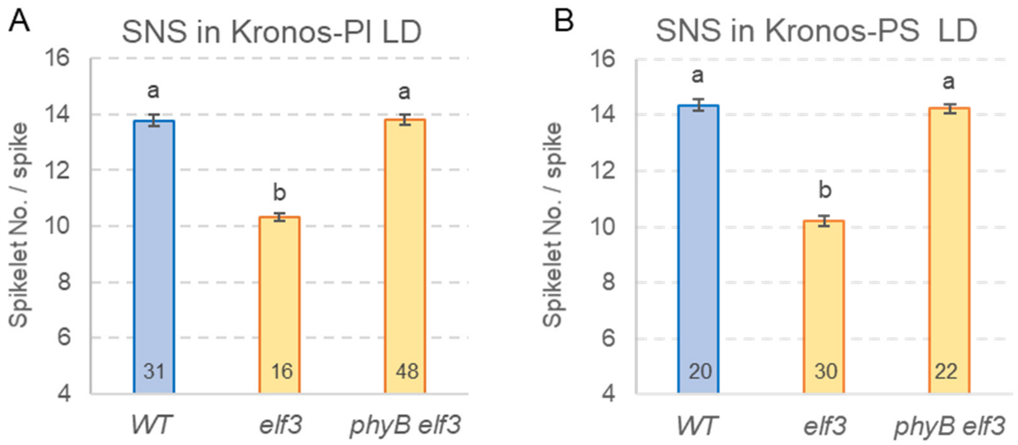

**S3 Fig. Transcript levels of flowering genes *VRN1*, *VRN2*, *CO1*, and *CO2* in Kronos PI, *phyB* and *elf3 phyB*.** (**A-B**) *VRN1*, (**C-D**) *VRN2*, (**E-F**) *CO1*, and (**G-H**) *CO2.* (**A**, **C**, **E**, and **G**) Wildtype *vs*. *phyB*. (**B**, **D**, **F**, and **H**) Wildtype *vs*. *elf3 phyB*. Primers used for qRT-PCR amplify both homoeologs of each gene. The WT data is the same within each row but can be at different scales. Error bars are s.e.m based on 5 biological replications. ns = not significant, * = $P < 0.05$, ** = $P < 0.01$, *** = $P < 0.001$ based on *t*-tests between mutants and wildtype at the different time points. Raw data and statistics are available in Data I in S1 Data.

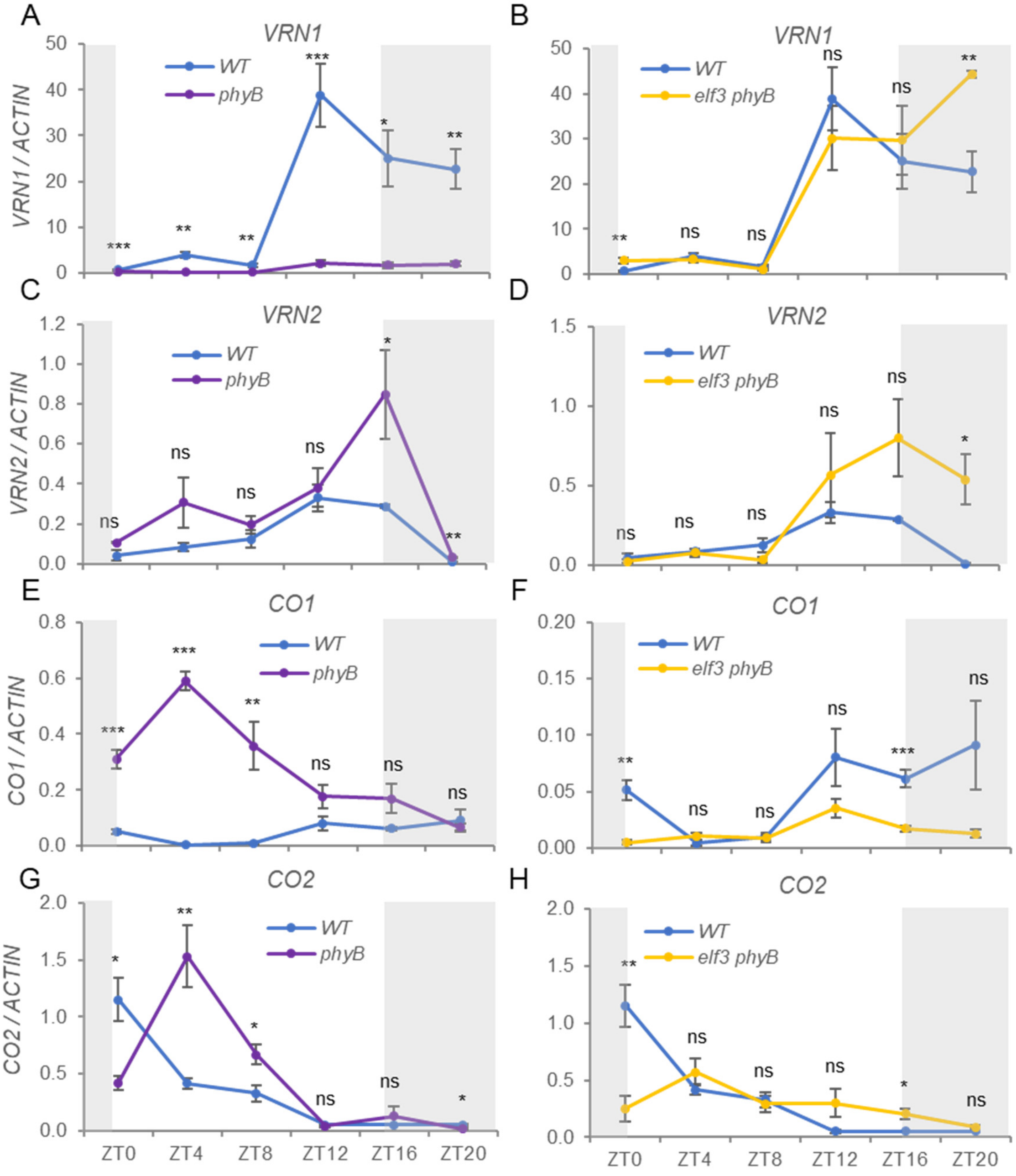

**S4 Fig. *ELF3* transcript levels in leaves in the presence of *phyB* and *phyC* mutants under different photoperiods.** (**A**) Transcript levels of *Elf3* during the day in PS under SD. (**B**) *Elf-A3* and *Elf-B3* transcripts per million (TPM) in PI WT, *phyB* and *phyC* in leaves collected at ZT4 from 4-w old plants grown under LD and 8-w old plants grown under SD. Data are from previously published RNA-seq (Kippes et al. 2020). (**C**) Transcript levels during the day in PI and *phyC* under LD (**D**) Transcript levels during the day in PI and *phyB* under LD. Transcript levels were determined by qRT-PCR using *ACTIN* as endogenous control. NS = $P > 0.05$. Raw data and statistics are in Data J in S1 Data.

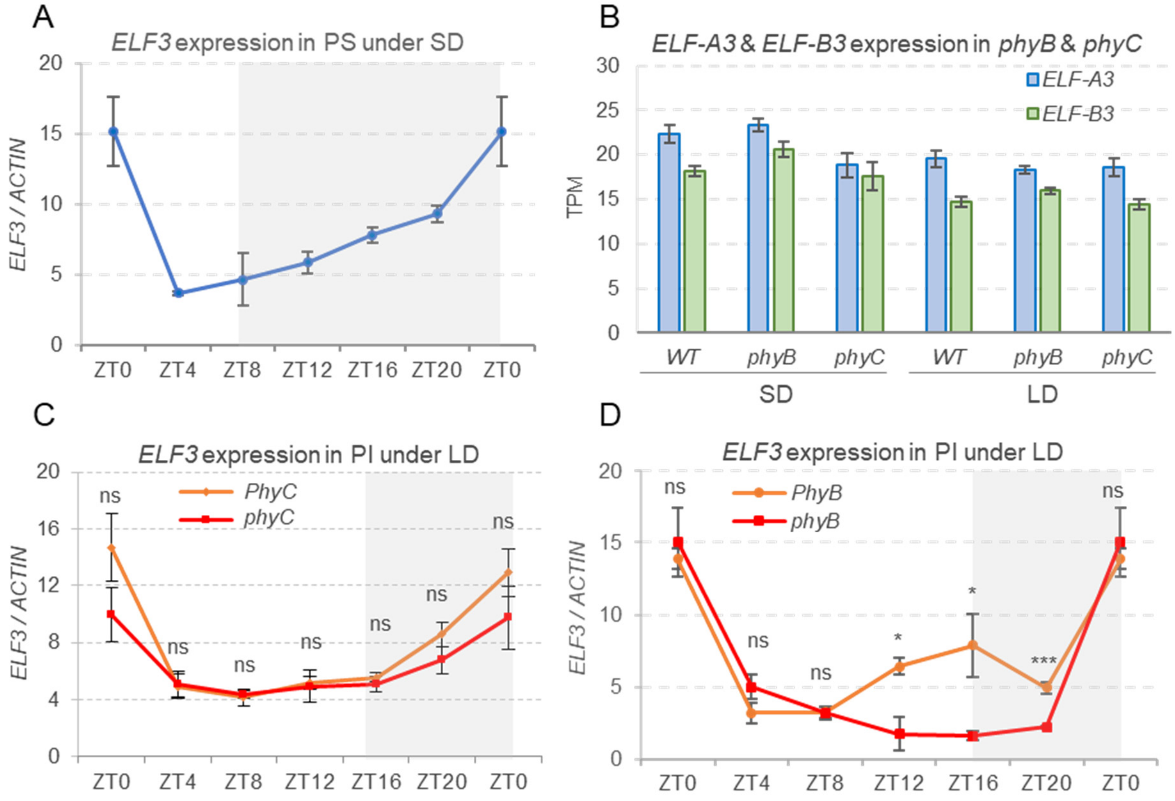

**S5 Fig. Effect of the constitutive expression of *ELF3* under the maize *UBIQUITIN* promoter (UBI::ELF3:HA).** (**A-C**) Transcription profiles in the 7th leaf collected at ZT0, ZT4, ZT12 and ZT20 from five-week-old Kronos PS plants grown under LD. *ACTIN* was used as endogenous control. (**A**) *ELF3*, (**B**) *PPD1* (conserved primers that amplify both *Ppd-A1b* and *Ppd-B1b*) and (**C**) *FT1*. Different letters indicate significant differences in Tukey tests ($P < 0.05$). The colors of the letters match the color of the respective treatment. OE = UBI::ELF3:HA and NT= non-transgenic sister line. (**D**) Complementation of the *elf3* mutant by UBI::ELF3:HA. A factorial ANOVA showed highly significant effects for both the mutant and transgenic genotypes and for their interaction. Simple effects were tested by contrasts: ns = not significant, ** = $P < 0.01$, *** = $P < 0.001$. Raw data and statistics are available in Data K in S1 Data.

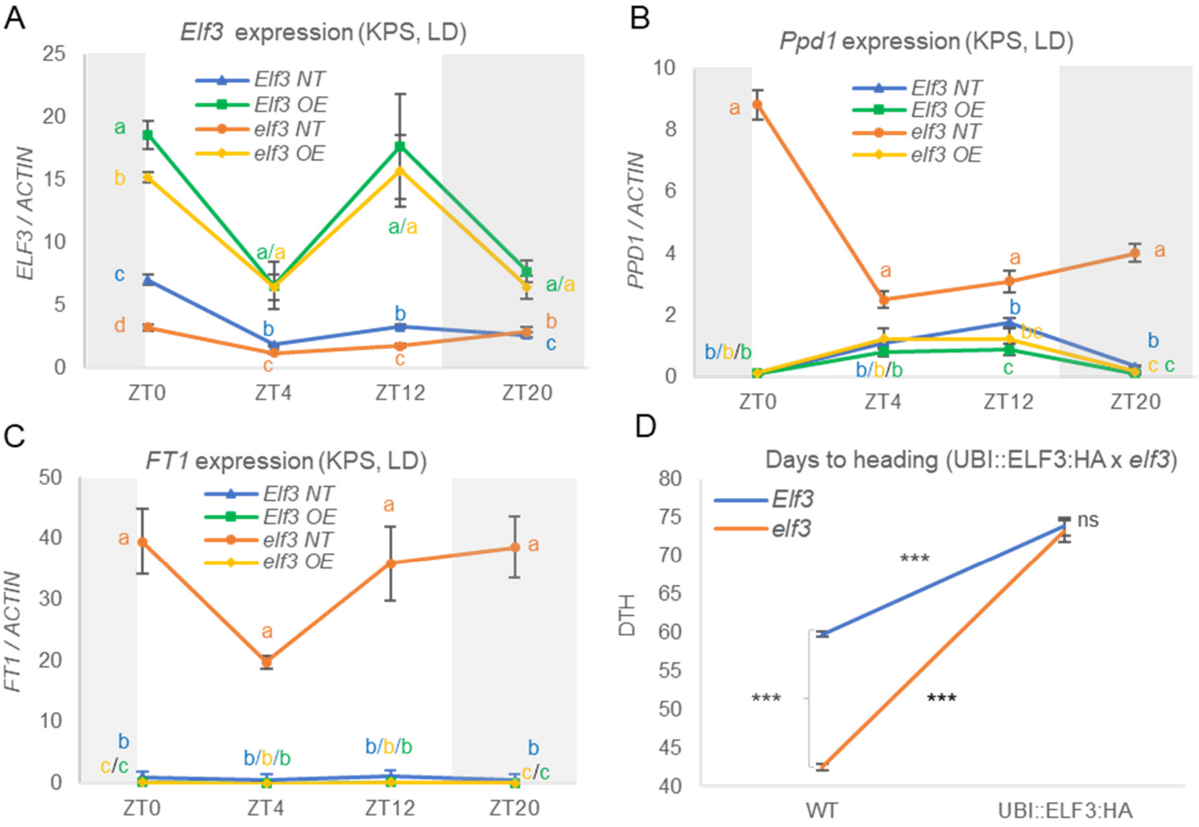

**S1 Table**. EMS-induced nucleotide changes that resulted in premature stop codon mutations in *ELF3*, *PHYB*, and *PPD-A1*. The ID of each of the mutant lines, the position of the loss-of-function mutations in the CDS and protein, and the primers used in the diagnostic KASP assays are listed. Capital letters in the primer sequences indicate the VIC and FAM tails. The 3′ allele-specific nucleotides are underlined.

| Gene | Mutant | Mutation | Effect | Primer Name | Primer Sequence (5' to 3') |
|---|---|---|---|---|---|
| *Elf-A3* | T4-928 | G1596A | W532* | ELFA3-SP-F1 | GCTTCACCATCTCAAGACAATGAT |
| | | | | ELFA3-WT-R1 | GAAGGTGACCAAGTTCATGCTtgaggcggaggagcacac |
| | | | | ELFA3-MUT-R2 | GAAGGTCGGAGTCAACGGATTtgaggcggaggagcacat |
| *Elf-B3* | T4-3669 | C1576T | Q529* | ELFB3-SP-F1 | CTTCACCATCTCAAGACAATGAC |
| | | | | ELFB3-WT-R1 | GAAGGTGACCAAGTTCATGCTggagcacaccagttgttctg |
| | | | | ELFB3-MUT-R2 | GAAGGTCGGAGTCAACGGATTggagcacaccagttgttcta |
| *PhyB-A* | T4-2711 | C1756T | R586* | PHYB-A-SP-F1 | TCACGCCGAGTGATTATTTG |
| | | | | PHYB-A-WT-R1 | GAAGGTGACCAAGTTCATGCTaaatgccttgaatgatgatcg |
| | | | | PHYB-A-MUT-R2 | GAAGGTCGGAGTCAACGGATTgaaatgccttgaatgatgatca |
| *PhyB-B* | T4-2078 | C3079T | Q1027* | PHYB-B-SP-F1 | TTTTCACTTGGAAATGTTATGAATGC |
| | | | | PHYB-B-WT-R1 | GAAGGTGACCAAGTTCATGCTtcatcagggatatctcgaataagttg |
| | | | | PHYB-B-MUT-R2 | GAAGGTCGGAGTCAACGGATTtcatcagggatatctcgaataagtta |
| *Ppd-A1* | T4-689 | G671A | W154* | PPDA1-WT-F1 | GAAGGTGACCAAGTTCATGCTgaacgagcttaagaacctctgg |
| | | | | PPDA1-MUT-F2 | GAAGGTCGGAGTCAACGGATTagaacgagcttaagaacctctga |
| | | | | PPDA1-SP-R1 | ACTACTGTGGAAGAAGAGAACAGAC |

**S2 Table**. Primers used for qRT-PCR analysis.

| Gene | Forward Primer (5' to 3') | Reverse Primer (5' to 3') | Reference |
|---|---|---|---|
| *ACTIN* | ACCTTCAGTTGCCCAGCAAT | CAGAGTCGAGCACAATACCAGTTG | Distelfeld et al. (2009) |
| *CO1* | CACATCAGAGTGGTTATGC | GGACTGGACCGTATTGTC | Chen et al. (2014) |
| *CO2* | AAGGGTGTGAGTGTGTAG | GATATGTCATTGCTGATGGAAG | Chen et al. (2014) |
| *FT1* | CAGCAGCCCAGGGTTGAG | ATCTGGGTCTACCATCACGAGTG | Yan et al. (2006) |
| *Ppd-A1* | AGACAAGGCTGATGAAACGA | CGATGGATTGACCAAACTG | Shaw et al. (2012) |
| *Ppd-B1* | AAGACAAGGTTGATGACGTGA | GAGGGATTGATCACGTTGG | Shaw et al. (2012) |
| *Vrn1* | AAGAAGGAGAGGTCACTGCAGG | GGCTGCACTGCCGCA | Yan et al. (2006) |
| *Vrn2* | CCACCATCGTGCCATTCT | CCCACCATCATCTCTGTATCAA | Distelfeld et al. (2009) |

**S3 Table**. Primers used for cloning ELF3 and PHYB in yeast-two-hybrid assays.

| Gene | **Primer (5' to 3') [a]** |
|---|---|
| ELF3-GW-F | GGGGACAAGTTTGTACAAAAAAGCTGCCACCATGAGGAGGGCCGGCGGC |
| ELF3-GW-R | GGGGACCACTTTGTACAAGAAAGCTGAACGCGGGCCGTTCTGCTGCCT |
| N-PHYB-F | CC**GAATTC**ATGGCCTCGGGAAGCCGCGC |
| N-PHYB-R | TCCC**CCCGGG**TGCATCTCTGAAGGAGTCCCG |
| C-PHYB-F | CC**GAATTC**GGAGAGGGCACTAGTAACTC |
| C-PHYB-R | CC**ATCGAT**GCTCCGATCCCTACTTTCT |

[a] Underlined indicates attB sites for gateway recombination. Bold indicates restriction sites

**S4 Table**. Primers used for chromatin immunoprecipitation.

| Amplified segment | Primer | Coordinates from start |
|---|---|---|
| PPD1 CDS +1514_F | AAGACAAGGTTGATGACGTGA | +1514 |
| PPD1 CDS +1514_R | GAGGGATTGATCACGTTGG | +1904 |
| PPD1-142_CHIP_F | GATGCGACCGAGGTTCGA | -142 |
| PPD1-142_CHIP_R | GCCTGACTCCAAGAGGAAACATG | -21 |
| PPD1-446_CHIP_F | GCTCTGTTCCTGCCCGATTG | -446 |
| PPD1-446_CHIP_R | CTCCAGCAATTTCCGGGCAC | -320 |
| PPD1-460_CHIP_F | CCTGTCTGTCACTCGTCTGC | -460 |
| PPD1-460_CHIP_R | GGTTAATCTCCAGCGATTTC | -313 |
| PPD1-983_CHIP_F | TGATTGAGAGGCGAGCGAGG | -983 |
| PPD1-983_CHIP_R | CGCGCTGGATCCGCATATCT | -796 |
| FUL2 -1933_ CHIP_F | ACCTCGAGGTCGAGATGCAGTAC | -1933 |
| FUL2 -1933_ CHIP_R | TAGACCTATCACCGGCGCGT | -1750 |

### References


Alvarez MA, Tranquilli G, Lewis S, Kippes N, Dubcovsky J. Genetic and physical mapping of the earliness *per se* locus *Eps-A*$^{m}$*1* in *Triticum monococcum* identifies *EARLY FLOWERING 3* (*ELF3*) as a candidate gene. Funct Integr Genomic. 2016;16:365-82.

Beales J, Turner A, Griffiths S, Snape JW, Laurie DA. A pseudo-response regulator is misexpressed in the photoperiod insensitive *Ppd-D1a* mutant of wheat (*Triticum aestivum* L.). Theor Appl Genet. 2007;115(5):721-33.

Chen A, Li C, Hu W, Lau ML, Lin H, Rockwell NC, Martin SS, Jernstedt JA, Lagarias JC, Dubcovsky J. *PHYTOCHROME C* plays a major role in the acceleration of wheat flowering under long day photoperiod. Proc Natl Acad Sci USA. 2014;111:10037-10044.

Distelfeld A, Tranquilli G, Li C, Yan L, Dubcovsky J. Genetic and molecular characterization of the *VRN2* loci in tetraploid wheat. Plant Physiol. 2009;149:245-257.

Pearce S, Kippes N, Chen A, Debernardi JM, Dubcovsky J. RNA-seq studies using wheat *PHYTOCHROME B* and *PHYTOCHROME C* mutants reveal shared and specific functions in the regulation of flowering and shade-avoidance pathways. BMC Plant Biol. 2016;16:141.

Pearce S, Shaw LM, Lin H, Cotter JD, Li C, Dubcovsky J. Night-break experiments shed light on the *Photoperiod1*-mediated flowering. Plant Physiol. 2017;174:1139-50.

Shaw LM, Turner AS, Laurie DA. The impact of photoperiod insensitive *Ppd-1a* mutations on the photoperiod pathway across the three genomes of hexaploid wheat (*Triticum aestivum*). Plant J. 2012; 71:71–84.

Wilhelm EP, Turner AS, Laurie DA. Photoperiod insensitive *Ppd-A1a* mutations in tetraploid wheat (*Triticum durum* Desf.). Theor Appl Genet. 2009;118:285-94.

Yan L, Fu D, Li C, Blechl A, Tranquilli G, Bonafede M, Sanchez A, Valarik M, Yasuda S, Dubcovsky J. The wheat and barley vernalization gene *VRN3* is an ortholog of *FT*. Proc Natl Acad Sci USA. 2006;103:19581-19586.
